# Adipose-Tumor Crosstalk contributes to CXCL5 Mediated Immune Evasion in PDAC

**DOI:** 10.1101/2023.08.15.553432

**Authors:** R. McKinnon Walsh, Joseph Ambrose, Jarrid L. Jack, Austin E. Eades, Bailey Bye, Mariana T. Ruckert, Appolinaire A. Olou, Fanuel Messaggio, Prabhakar Chalise, Dong Pei, Michael N. VanSaun

## Abstract

**Background:** CXCR1/2 inhibitors are being implemented with immunotherapies in PDAC clinical trials. Cytokines responsible for stimulating these receptors include CXCL ligands, typically secreted by activated immune cells, fibroblasts, and even adipocytes. Obesity has been linked to poor patient outcome and altered anti-tumor immunity. Adipose-derived cytokines and chemokines have been implicated as potential drivers of tumor cell immune evasion, suggesting a possibility of susceptibility to targeting specifically in the context of obesity.

**Methods:** RNA-sequencing of human PDAC cell lines was used to assess differential influences on the cancer cell transcriptome after treatment with conditioned media from peri-pancreatic adipose tissue of lean and obese PDAC patients. The adipose-induced secretome of PDAC cells was then assessed by cytokine arrays and ELISAs. Lentiviral transduction and CRISPR-Cas9 was used to knock out CXCL5 from a murine PDAC cell line for orthotopic tumor studies in diet-induced obese, syngeneic mice. Flow cytometry was used to define the immune profiles of tumors. Anti-PD-1 immune checkpoint blockade therapy was administered to alleviate T cell exhaustion and invoke an immune response, while the mice were monitored at endpoint for differences in tumor size.

**Results:** The chemokine CXCL5 was secreted in response to stimulation of PDAC cells with human adipose conditioned media (hAT-CM). PDAC CXCL5 secretion was induced by either IL-1β or TNF, but neutralization of both was required to limit secretion. Ablation of CXCL5 from tumors promoted an immune phenotype susceptible to PD-1 inhibitor therapy. While application of anti-PD-1 treatment to control tumors failed to alter tumor growth, knockout CXCL5 tumors were diminished.

**Conclusions:** In summary, our findings show that known adipokines TNF and IL-1β can stimulate CXCL5 release from PDAC cells *in vitro. In vivo*, CXCL5 depletion alone is sufficient to promote T cell infiltration into tumors in an obese setting, but requires checkpoint blockade inhibition to alleviate tumor burden.

**DATA AVAILABILITY STATEMENT:** Raw and processed RNAseq data **will be** further described in the GEO accession database (**awaiting approval from GEO for PRJ number**). Additional raw data is included in the supplemental material and available upon reasonable request.

**WHAT IS ALREADY KNOWN ON THIS TOPIC:** Obesity is linked to a worsened patient outcome and immunogenic tumor profile in PDAC. CXCR1/2 inhibitors have begun to be implemented in combination with immune checkpoint blockade therapies to promote T cell infiltration under the premise of targeting the myeloid rich TME.

**WHAT THIS STUDY ADDS:** Using *in vitro/ex vivo* cell and tissue culture-based assays with *in vivo* mouse models we have identified that adipose derived IL-1β and TNF can promote tumor secretion of CXCL5 which acts as a critical deterrent to CD8 T cell tumor infiltration, but loss of CXCL5 also leads to a more immune suppressive myeloid profile.

**HOW THIS STUDY MIGHT AFFECT RESEARCH, PRACTICE, OR POLICY:** This study highlights a mechanism and emphasizes the efficacy of single CXCR1/2 ligand targeting that could be beneficial to overcoming tumor immune-evasion even in the obese PDAC patient population.

## BACKGROUND

Pancreatic cancer is projected to become the second leading cause of cancer associated death in the United States by 2025, despite being projected to be eleventh in incidence, with Pancreatic ductal adenocarcinoma (PDAC) incidence constitutes ∼90% of cases (1). Obesity has been implicated as a risk factor enhancing tumor progression and metastatic potential of PDAC, among other cancers (2–6), and high body mass index (BMI) correlates with earlier incidence and poorer outcome in retrospective studies of PDAC (7,8). Obesity leads to the induction inflammation in adipose tissue: thus termed “reactive adipose” resulting in a consequential altered secretion of adipose derived cytokines, or adipokines (9,10). The direct influence of adipokines on intrinsic cancer cell pro-tumorigenic properties such as proliferative capacity and metastatic potential have been well documented in a variety of contexts (2,11,12); secondary effects of adipokines that alter the tumor’s relationship with other cells in the tumor microenvironment members as well as systemic effects, however, are only more recently being intimately investigated (13). Adipose derived IL-1β has been shown to be responsible for preventing the infiltration of CD8 T cells in obese mouse models of PDAC (4), and that by blocking IL-1β this infiltration can be rescued and prevents infiltration of a portion of monocyte lineage immune cells (CD11b/Ly6G+). Because T cell infiltration into PDAC tumors is often poor, checkpoint blockade inhibitors have thus far been ineffective in pre-clinical models or clinical trials without additional perturbation of the tumor or microenvironment (14–17). Therefore, determining potential mechanisms of enhancing T cell infiltration is recognized as an utmost important achievement in PDAC.

Of the successful inductions of immunotherapy sensitivity, many studies have focused toward affecting the myeloid immune profile through the blockade of CXCR1/2 and its ligands which are oft over-expressed in PDAC (18–24) Currently, clinical trials are testing the efficacy of CXCR1/2 inhibition in combination with immune checkpoint blockade (ICB) therapy. Multiple cytokines and chemokines that target these receptors have been identified as upregulated in the context of PDAC(24), yet it was only high CXCL5 expression that displayed significantly worsened patient outcome in TCGA-PAAD datasets (Supplemental Figure 2). CXCL5 has been identified as another CXCR2 ligand that drives neutrophil recruitment and activation (25), and has been associated with poor patient outcome (19,26). Still, pre-clinical studies have shown impressive results in targeting other individual CXCR1/2 ligands such as CXCL1 (18) or IL-8 (22,27).

In this study, we show that tumor secretion of chemokine CXCL5 can be directly induced by IL-1β or Tumor Necrosis Factor (TNF), which can be sourced in high levels from adipose tissue. We then used a combination of InVivoMAb anti-mouse PD-1 (CD279) [BE0273, Clone 29F.1A12] combined with a depletion of CXCL5 by Crispr-Cas9 mediated knockout to interrogate the contribution of CXCL5 to susceptibility for immune-checkpoint blockade therapy in the context of obesity.

## METHODS

### Cell culture

Human cell lines MiaPaCa2, AsPC1, CaPan2, HPNE-G12D, Panc1, HEK-293T, adMSC, and 3T3-L1 cell lines were acquired from ATCC and used within 15 passages from purchase.

The K8484 cell line was a kind gift from Dr. Dave Tuveson, Cold Spring Harbor, New York, USA harboring a p53^R172H/+^ and Kras^G12D/+^ mutation driven by Pdx1^Cre^, (28). Cells were grown in complete Dubelco’s modified eagle media (DMEM) high-glucose, pyruvate [Gibco™ 11995073], supplemented with Antibiotic-Antimycotic (Anti-Anti) [Gibco™ 15240062] and 5-10% Fetal Bovine Serum [Biowest S1620] at 37°C in 5% CO2 [Matheson]. All cell lines were routinely monitored for mycoplasma. Murine cell lines were authenticated for epithelial or fibroblastic markers by western blot or flow cytometry.

### Patient-derived human adipose tissue conditioned media (hAT-CM) collection

Patient samples were provided by the KUMC Biospecimen Repository Core Facility. Peri-pancreatic adipose tissue was collected at time of surgical resection of tumor and kept at room temperature in PBS until processing. Gross peri-pancreatic adipose tissue was washed in 3x in room temperature PBS + Anti-Anti + 10 μg/mL Ciprofloxacin [Santa Cruz 29064], then cut and weighed into small sections. Adipose was incubated for 1hr in 5-10 mL of serum-free DMEM media supplemented with Anti-Anti and Ciprofloxacin to clear any factors released due to manipulation (29). Then, 1 mL of serum-free DMEM + Anti-Anti + Ciprofloxacin was added per 125 mg of adipose tissue in a 6 well plate and incubated for 24h at 37°C 5% CO2 [Matheson]. Conditioned media was collected using a filtered 1 mL pipette, then spun at 300 x g for 5 minutes to remove cells, then spun at 5000 x *g* for 5 minutes to further remove other debris. The media was drawn up in a Luer-Lock Syringe [BD 309628, BD 309646] then sterile filtered with a 0.22 μM syringe filter [Rephile RJP3222NH], aliquoted to 1.7 mL tubes, and stored at -80C until use application to cancer cells or for enzyme-linked immunosorbent assay. hAT-CM was applied to cells for treatment at a ratio of 1:3-1:5 hAT-CM:Serum Free DMEM.

### Proliferation assays

Proliferation of Cas9-NTC (non-targeting control) or Cas9-CXCL5 K8484 cells was measured by EdU incorporation as previously described (30). Cells were plated at 70k/well on a 24 well plate (K8484 cells) or 10k/well on a 96 well plate (conditioned media experiments). Cells were treated for 24h with drug or conditioned media (1:3 dilution hAT-CM:SF DMEM). EdU was diluted to 1 mM in SF DMEM added to the wells at 10 μM final concentration, mixed, and cells were incubated for an additional 6h with EdU prior to stopping the assay. Processing of EdU incorporated cells was performed as previously described (30). Briefly, cells were washed with ice cold PBS 2x, trypsinized into single cell suspension, and fixed overnight in 1.5% Buffered Formalin rocking at 4C. Cells were washed, permeabilized, and underwent Click-It reaction using CuSO_4_ (Alfa Aesar 14178) and 647 Azide-Dye (Invitrogen A10277), counter-stained with propidium iodide (ThermoFisher P3566) and measured for fluorescence using an Attune NxT Flow Cytometer.

### RNA-seq sample preparation

Cell lines were trypsinized using 0.25% Trypsin EDTA [Gibco™ 25200056] from 10 cm culture dishes and then neutralized with complete media once detached from the plate. Cells were counted and spun at 300 x *g* for 5 minutes, then resuspended in complete media and distributed at 1E6 per well in a 6 well plate. Cells were allowed to settle overnight, washed 2x with serum free media, then treated with conditioned media diluted 1:3 using serum free DMEM + Anti-Anti as diluent. Cells were cultured for 24 hours, washed 2x with 1 mL of ice-cold PBS [Sera-Care 5460-0023], and then lysed in 300 uL QIA-zol [Qiagen 51-13-02]. RNA was extracted using Direct-zol RNA Miniprep Kit [Zymo Research R2052] following the manufacturer’s protocol.

The Library prep was completed following the Tecan Universal Plus mRNA library Prep (Stranded mRNA Library) and then sequenced on a NovaSeq S1 200 cycle (100PE) flow cell using XP 2-lane chemistry for independent lane loading. A second sequencing run of the first set was used to provide supplemental reads for samples HPNE_297 & HPNE_415. A second set of experiments were combined as more patient samples were accrued. Groupings of samples were MiaPaCa2, Panc1, SW1990, and HPNE-G12D.

There were slight differences in the gene expressions between the two batches of samples. Those batch effects were corrected using the ComBat-seq method (31) prior to carrying out any statistical analysis (**Supplemental Figure 1)**.

### RNAseq data analysis

For each RNA-seq Sample, QC was conducted by using fastqc (http://www.bioinformatics.babraham.ac.uk/projects/fastqc). Samples that passed QC were then aligned to the human genome (hg38) with RSEM (32) (PMID: 21816040) and bowtie2 (33) (PMID: 22388286). Gene counts were used for downstream analysis. Prior to differential expression analysis, non-expressed genes (genes having very small or zero counts) were filtered out in order to remove the biologically uninteresting genes and also to attenuate the burden of multiple testing. Differential expression analyses of the gene expression data among the groups were carried out using a generalized linear model based on negative binomial distribution as implemented in R software package *edgeR*. Pairwise comparison of the gene expression between the groups were carried out by specifying appropriate contrast statements in the modeling framework. Multiple testing adjustments were carried out using Benjamini-Hochberg’s False Discovery Rate (FDR) method.

### RNA-seq Pathway Analysis

Pathway analysis was performed using Qiagen Ingenuity Pathway Analysis (IPA). Differentially expressed genes between lean and obese treated samples (obese as reference) were input with logFC cutoff of -.2 to .2 exclusion, and p-value minimum of .8. Upstream analysis of molecules was filtered for cytokines and growth factors. The top 25 activation-z-score cytokines and growth factors were input into SRplot (http://www.bioinformatics.com.cn/srplot) for visualization. Gene expression consistency percent was generated by dividing the number of genes with measurement direction consistent with the activation of the cytokine/growth factor by the total genes expected to be directionally changed by the cytokine/growth factor as predicted by IPA.

### Cytokine Array

Cytokine release from adipose stimulated MiaPaca2 cells were compared to unstimulated MiaPaca2 cells using a Raybio C-Series Human Cytokine Antibody Array C3 (AAH-CYT-3). The protocol was followed according to the manufacturer’s recommendations. Conditioned media was collected from human adipose and then applied to MiaPaca2 cells, subsequently washed and then these stimulated cells were cultured in serum free media for an additional 24h to collect tumor cell conditioned media. Undiluted culture media was loaded onto the array for analysis.

### Enzyme Linked Immunosorbent Assays (ELISAs)

CXCL5 ELISAs (Figure 1) samples were collected from our cancer cell line panel after plating the cells at 50k/well and allowed to settle overnight. The cells were washed and treated with 1:3 diluted hAT-CM for 36h based on timring from a previous study (34). Blocking assay (Figure 3): Panc1 cells were plated at 60k/well on a 48 well plate and allowed to settle overnight. The next day, cells were stained with LumiNuc LUCS 13 dye [LumiProbe 24010] at 2.5 uM for 1h in 2.5% FBS media at 37°C prior to washing 2x with serum free media for conditioned media treatment. Conditioned media/recombinant protein treatment of cells for CXCL5 release was performed similarly to preparation for RNA-seq analysis (1:4 dilution of hAT-CM:Serum Free media), but media was additionally supplemented with 1.5% BSA [Rockland BSA-30] for stabilization of peptides and allowed to release CXCL5 for 36h based on timing results from a previous study (34). The collected samples were transferred to a 96 well plate and spun at 1k x *g* to remove cells and debris after collection. The remaining cells were fixed in formalin and imaged on a Sartorius Incucyte to obtain nuclear event count. The collected media was then measured for CXCL5 using a Human CXCL5/ENA-78 DuoSet ELISA kit [R&D Systems DY254] and DuoSet ELISA Ancillary Reagent Kit 2 [R&D Systems DY008B] according to the manufacturer’s protocol. Due to the limited availability of tissue samples, analyses were performed by pooling samples of adipose conditioned media to stimulate cells. Recombinant human TNF [PeproTech 300-01A] and IL-1β [PeproTech 200-01B] were applied at 1 μg/mL. Blocking antibodies against TNF [CST 7321S] and IL-1β [abcam ab9722] or IgG [Santa Cruz sc-2027] were applied at 1 ug/mL. TNF ELISA [R&D Systems DY210-05] and IL-1β ELISA [R&D Systems DY201-05] were performed according to the manufacturer’s protocols with Ancillary Reagent Kit 2 [R&D Systems DY008B].

**Figure 1:**
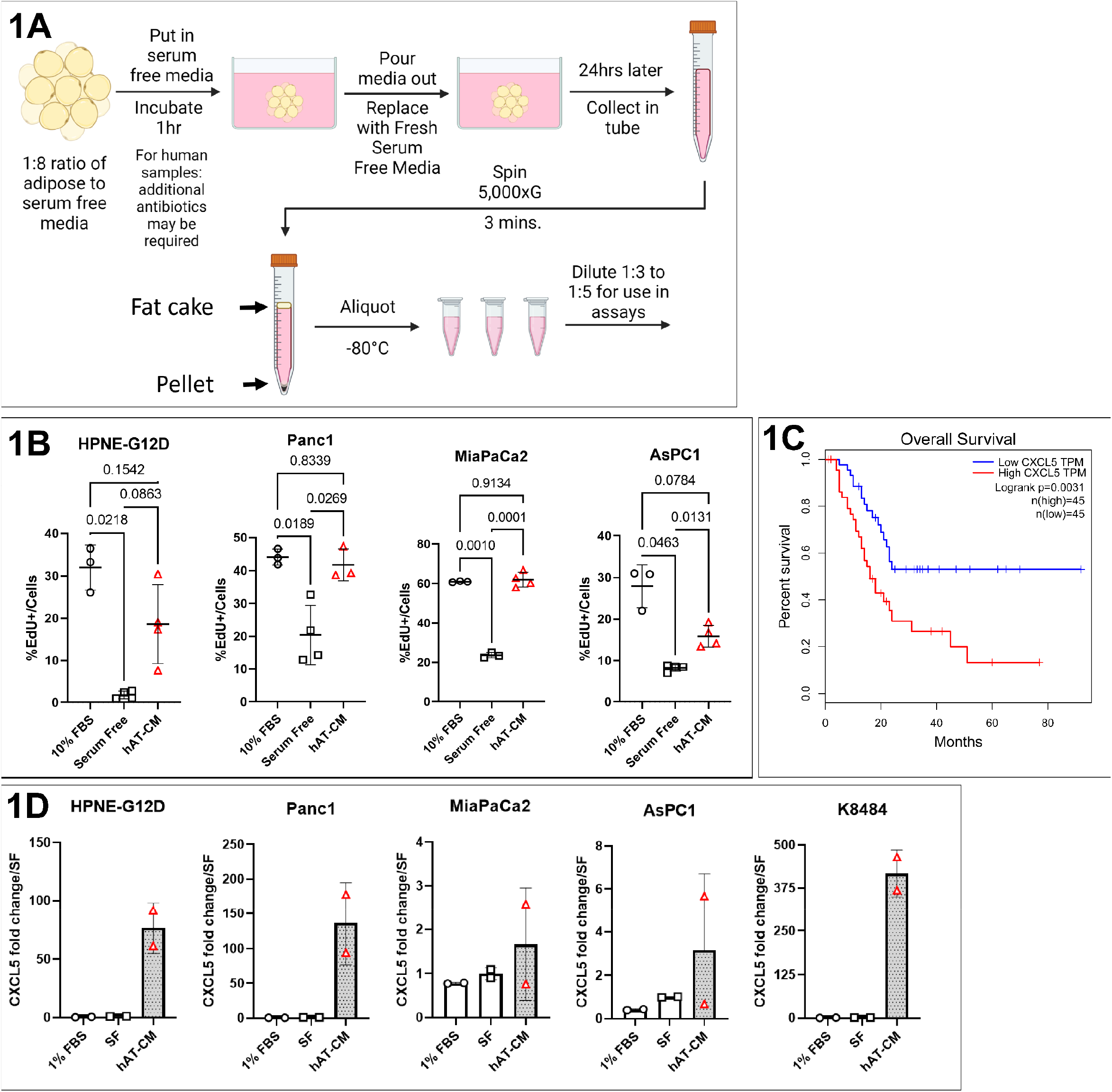
Figure 1A: Schematic created in BioRender illustrating human adipose tissue conditioned media (hAT-CM). Figure 1B: Proliferation measured by EdU incorporation of cells *in vitro* in response to 24h of stimulus with pooled hAT-CM. Figure 1C: Kaplan-Meier Plot generated using GEPIA showing significant differences between the upper and lower quartile expressing patients in the PAAD-TCGA dataset. Figure 1D: Stimulation of cells with hAT-CM shows CXCL5 in the media of the stimulated cells measured by ELISA. Statistical analyses were performed using Brown-Forsythe and Welch corrected One-Way Anova in GraphPad Prism.

### Lentivirus Production

HEK-293T (ATCC) cells were treated according to the Lipofectamine 3000™ protocol with Opti-MEM [Gibco™ 31985070] and Lipofectamine™ [Invitrogen L3000001] + plasmid DNA [Lenti-Cas9 Addgene #52962; Lenti-envelope pCMV-VSV-G Addgene #8454; Lenti-backbone pCMV-dR8.2 dvpr Addgene #8455] to produce virus containing media. The lentivirus was concentrated with Lenti-X™ concentrator [TakaraBio 631231].

### CRISPR/Cas9 knockout of CXCL5

TransOMIC technologies CCMS1001 kit was used for generation of guide RNA for knockout studies. Two separate guide RNAs [TransOMIC technologies TEVM-1096303 (“3”), TEVM-1163445 (“45”)] were selected and used to generate the four CXCL5-knockout clones used for experiments. Sequences are as follows: NTC (GGAGCGCACCATCTTCTTCA), “3” (ATGGCGAGATGGAACCGCTG), “5” (GTTCCATCTCGCCATTCATG). K8484 (28) cells were transfected with a lenti-viral, blasticidin resistant Cas9 expression cassette [Addgene 52962], then transfected with a lenti-viral CXCL5 guide RNA with puromycin resistance and maintained in media supplemented with 10 μg/mL Puromycin [Gibco™ A1113803], and plated at single cell level on a 96 well plate. Two clones from each targeting guide were selected based on qPCR expression of CXCL5 (Qiagen QT01658146) and used for assays within 10 passages of selection.

### qPCR and quantification

qPCR was performed according to the three-step method in the protocol from APExBIO HotStart [APExBIO K1070] in a ViiA 7 Real-Time PCR System (ThermoFisher). initial denaturation (hold; 1 cycle): 95°C for 2 min; 40 cycles of denaturation (95°C for 15s), annealing (50–60°C for 30s) and extension (72°C for 30s) followed by 1 cycle of 95°C for 15s, 60 °C for 1 min and 95 °C for 15s. Quantification was performed by the Livak method (35) with GAPDH as the reference gene.

CXCL5 primer (Qiagen Mm_Cxcl5_2_SG QuantiTect Primer Assay QT01658146) GAPDH primer (Qiagen Mm_Gapdh_3_SG QuantiTect Primer Assay QT01658692)

### Animal Studies

All animal studies were performed in accordance with the approved IACUC Protocols at the University of Kansas Medical Center.

### Diet-induced Obesity (DIO)

Jax/BL6 mice at 6-8 weeks of age were put on a Western style High Fat Diet (HFD) [Teklad-Envigo InotivCo TD.88137], Adjusted Calories Diet (42% kcal from fat, 34% sucrose by weight) for a minimum of 3 months and maintained on HFD through experiment endpoint.

### Orthotopic Pancreatic Tumor Model

Cells were trypsinized, washed in PBS, counted, and 2E6 cells were pelleted at 1k x *g* for 3 minutes and kept on ice until resuspension. Mice were anesthetized using 3-5% Isoflurane [Patterson 14043070406] via Bickford Isoflurane Vaporizer and placed ventral side up on a sterile drape atop a warming pad. Mice are pre-operatively administered 50 uL of Buprenorphine ER-LAB 0.5 mg/mL [ZooPharm] and 300-500 uL of sterile 0.9% Sodium Chloride [Fresenius Kabi 918620] subcutaneously. Mice eyes were administered artificial tears [Patterson 14043-012-35] with sterile cotton tip applicators [Puritan 25-803]. The surgery site was cleared of fur using Nair for 5 minutes and wiped away with clean, dry or dampened paper towels. The surgery site was sterilized by wiping the skin with Povidone-Iodine Prep Pads [Dynarex 1108] and Isopropyl Alcohol Wipes [Medi-First 22133]. A paramedian incision is made in the abdomen and the pancreas is externalized. Pelleted cells were resuspended in 100 uL of Geltrex™ LDEV-Free Reduced Growth Factor Basement Membrane Matrix [Gibco™ A1413202]. 50 uL of cell/Geltrex™ solution was injected into the pancreas of syngeneic C57BL6 DIO mice using a 1 mL Slip Tip syringe [BD 329654] and a 27G x ½ inch needle [BD 305109]. Injections were monitored for leakage or proper delivery for 30 seconds, and the organ returned to its anatomical position. The peritoneum was sutured with 5-0 DemeCRYL™ [DemeTECH G175016F13M] and the skin was closed with a FST AutoClip® System [Fine Science Tools 12020-09, 12022-09]. The wound site was treated with triple antibiotic ointment [Akorn]. The wound clips were loosened on Day 6-7 and removed Day 7-8.

### Tumor dissociation for immune cell isolation

Mice were euthanized according to our IACUC approved protocols (2019-2493 and 22-02-226). The pancreas/tumors were removed from the mice, and a cross section was fixed in 10% buffered formalin overnight at room temperature, small chunks of tumor were frozen at -80C, and the remainder of the tumor was cleaned of normal pancreas for flow cytometry. The tumor was minced using a razor blade [FisherSci 12-640], resuspended in 10 mL of serum free RPMI-1640 [Gibco™ A1049101] supplemented with 1 mg/mL Soybean Trypsin Inhibitor (SBTI) [Sigma-Aldrich T6522-1G] in dH20. The resuspended tumors were spun at 350 x *g* 10 minutes at 4C, and resuspended in 5 mL of Tumor Digest Solution (see below) and shaken at 150-200 rpm for 25-35 minutes at 37C. 10 mL of 5% FBS RPMI-1640 was added to the digest solution, and the dissociated cells are filtered through a 100 μm cell strainer, and then a 40 μm strainer [Research Products International Corp 162427100, 162425100]. The original tube was washed with an additional 10 mL of RPMI Complete and put through the strainers to wash the remaining tissue. The dissociated tumors were then spun at 350 x *g* 20 minutes at 4C and resuspended in 1mL autoMACS® Running Buffer + 120U/mL DNase I.

Tumor Digest Solution (6 samples, 30 mL Prep): RPMI: 25mL, Collagenase P (15mg/mL stock, #11-213-857-001): 1mL (.5 mg/mL), DNase (30,000U/mL stock, DN25 Sigma in H2O): 120μL (120 U/mL), Collagenase V (2500CDU/mL, 20mg/mL stock in H2O [Sigma C9263]): 1mL (.66 mg/mL), SBTI (10mg/mL stock in H2O [Sigma-Aldrich T6522-1G]): 3mL (1 mg/mL)

### Spleen dissociation for immune cell isolation

Spleens were dissociated using a gentleMACS™ Dissociator [Miltenyi 130-093-235] in 3 mL of autoMACS® Running Buffer in gentleMACS™ C Tubes [Miltenyi 130-093-237] using the m_Spleen_01 protocol. The dissociated spleens were brought to 10 mL with autoMACS® buffer and spun at 350 x *g* for 10 minutes. The cells were resuspended in 10 mL of RBC Lysis Buffer [Biolegend 420301], vortexed, and incubated at RT for 10 minutes. The cells were spun again at 350 x *g* for 10 minutes and resuspended in 10 mL autoMACS® buffer + DNase I. The cells were then filtered through a 100 μm strainer, and spun at 350 x *g* 10 minutes at 4C and resuspended in 1 mL autoMACS® buffer.

### Cell Staining for Flow Cytometry

Our antibody panel is outlined in Supplemental Figure 4. Isolated cells (tumor and spleen) in 1 mL autoMACS buffer were transferred to 1.7 mL microfuge tubes and spun at 400 x *g* 10 minutes at 4C. Cells were then resuspended in 1 mL of 10 uL/mL TruStain FcX™ PLUS (anti mouse CD16/32) Antibody [Biolegend 156604] and blocked at RT for 20 minutes rocking. The cells were spun down and resuspended in 500 μL of autoMacs +DNase, and 125 μL was distributed to 3 other tubes for each of the immune panels. 125 uL of each of the antibody master mixes was distributed to each sample, briefly vortexed, and rocked at RT for 20 minutes in the dark. 700 μL of autoMACS + DNase I was added to the cells. The cells were spun down and resuspended in 1 mL autoMACS + DNase I twice more, and then resuspended in 300 μL of 50% Buffered Formalin:50% PBS and allowed to fix overnight at 4°C. The next day, cells were washed 2x by resuspending in 1 mL of autoMACS + DNAse I and spun at 600 x *g* for 10 minutes. The cells were finally resuspended into 300 μL of autoMACS and run on an LSR II [BD Biosciences] four-laser (405, 488, 552, 637 nm) analyzer. Compensation for controls were prepared using spleens or Ultracomp eBeads™ [Thermo Scientific 01-2222-42] as indicated in the supplemental data.

### *In vivo* PD1 administration

*InVivo*MAb anti-mouse PD-1 (CD279) [BioXCell #BE0273, Clone 29F.1A12] was prepared by dilution in *InVivo*Pure pH 7.0 Dilution Buffer (BioXCell #IP0070) to a working concentration of 200 μg/100 uL (2 mg/mL) treated with 100 μL (200 μg) per mouse (36) on days 10, 13, 16, 19, and 22 following tumor implantation. The antibody was delivered by intraperitoneal injection using a 28G ½ inch insulin syringe [BD 329461].

### Statistics

Statistical tests were performed as indicated in figure legends using GraphPad Prism 9.0: 1×1 comparisons were performed using Welch’s T-Test. Comparisons across multiple groups were carried out using One-way ANOVA with Browne-Forsythe and Welch corrections. Replicates are demoted in respective figure legends. TCGA data was accessed/Kaplan-Meier curves generated using GEPIA which provided statistical testing with figure generation (37).

## Results

### Adipose Conditioned Media from PDAC Patients promotes proliferation and CXCL5 secretion

In order to determine whether adipose factors directly influence pancreatic cancer cells we employed conditioned media transfer experiments. Peri-pancreatic adipose tissue was collected from pancreatic cancer patients during surgical resection and cultured for 24h-48h in serum-free media, then centrifuged to remove cells and debris (1A). The human adipose tissue conditioned media (hAT-CM) was diluted 1:3 in serum free media and applied to a series of primary and metastatic human PDAC cell lines, or an immortalized ductal cell line (HPNE-G12D) *in vitro* measuring proliferative capacity by EdU incorporation **Figure 1B**. hAT-CM treatment was sufficient to stimulate proliferation of all of the PDAC cell lines, but did not significantly increase EdU incorporation of the HPNE-G12D cell line above that of Serum Free media treatment.

Our next approach was taking media from MiaPaCa2 cells treated with pooled hAT-CM, and then assessing via cytokine array to identify changes to the secretome of the PDAC cells in response to the hAT-CM stimulation **(Supplemental Figure 2)**. CXCL5 was identified amongst differential potential targets when compared to untreated cells and selected for further evaluation across multiple human and murine PDAC cell lines by ELISA (**Figure 1C**). Of note, higher tumor CXCL5 expression is significantly correlated to poor patient outcome from TCGA data (19,26), while other CXCR1/2 ligands trend toward a correlative effect but failed to meet statistical significance (**Figure 1C, Supplemental Figure 2**).

### Differential IL-1β, TNF, and IFNy signaling are implicated as the pathway activators responsible for transcriptional changes in response to adipose factors

We hypothesized differential adipokine secretion sourced from obese versus lean adipose tissue could have contrasting effects when applied to PDAC cells. RNA-seq was performed on the pancreatic cell lines shown following treatment with hAT-CM collected from lean or obese patient adipose. Qiagen Ingenuity Pathway Analysis was used to process the data and identify potential upstream regulators driving transcriptional changes in PDAC and HPNE-G12D cells. IL-1β and TNF were identified as potential drivers of differential transcription between lean and obese hAT-CM treated cells (**Figure 2B**). Additionally, IFNγ signaling was implicated as a greater contributing factor by the RNA-seq upstream analysis displaying enhanced transcription or activation of associated target genes (**Supplemental Figure 3**). Heatmaps of individual samples across cell lines are shown with the top 500 differentially expressed genes (**Figure 2A**). Adipose media was assessed by ELISA for obese versus lean hAT-CM, which failed to show an expected (38) significant differential secretion of IL-1β and TNF (Supplemental **Figure 2D**), but did show large quantities being produced by the adipose tissue regardless of BMI.

**Figure 2:**
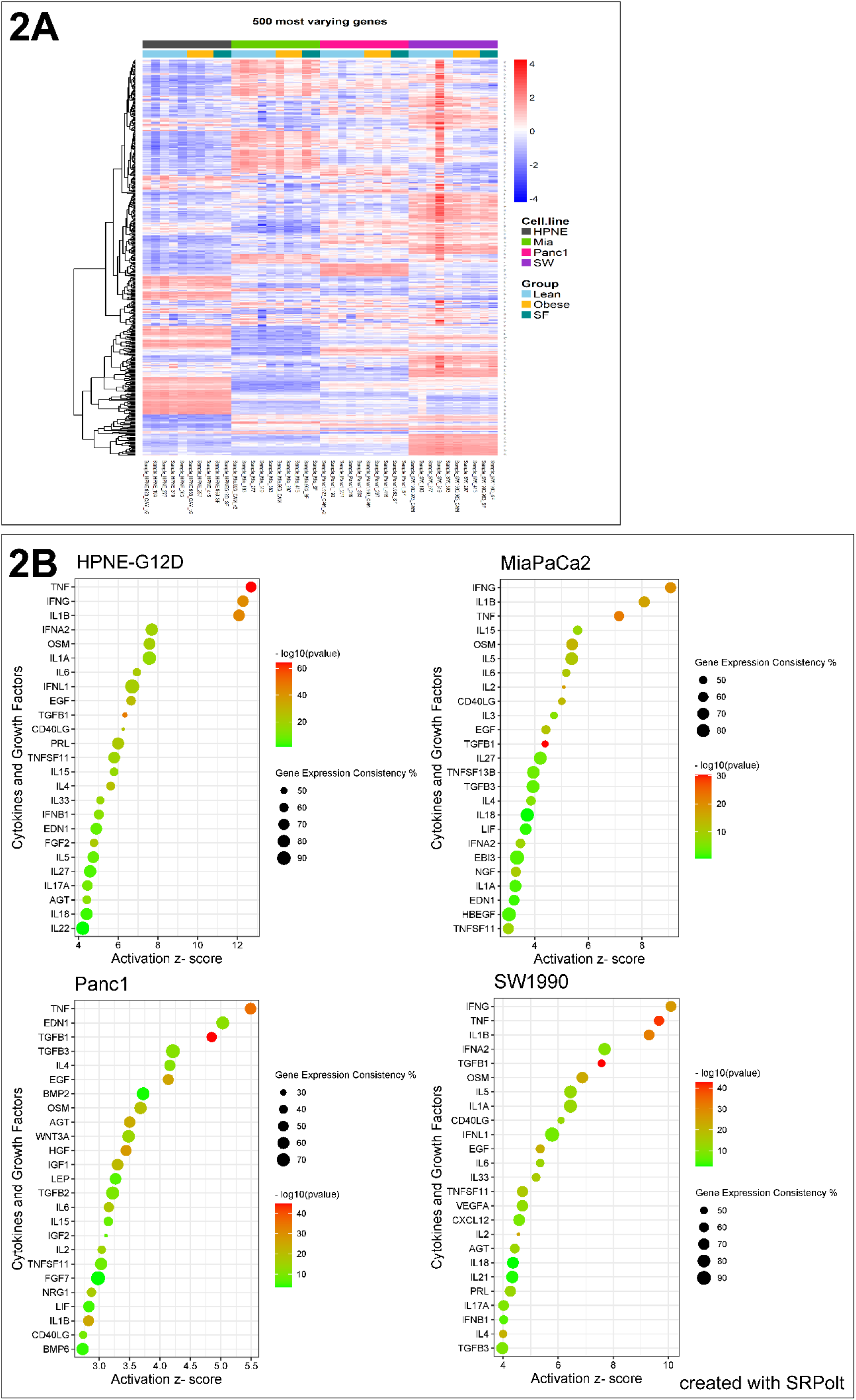
2A: Heatmap of top 500 differentially expressed genes across all groups. 2B: Bubble plots of top scored cytokines/growth factors in Ingenuity Pathway Analysis Upstream Analysis of Obese vs Lean RNA-seq data. Bubble plots were created by http://www.bioinformatics.com.cn/srplot, an online platform for data analysis and visualization.

### Blockade of adipose derived IL-1β and TNF diminishes induction of PDAC CXCL5 secretion

In order to evaluate the effect of hAT-CM associated IL-1β and TNF on PDAC cells, we applied recombinant ligands to stimulate CXCL5 secretion in culture. ELISA analysis of CXCL5 demonstrated that both recombinant IL-1β and TNF are capable of inducing CXCL5 secretion from PDAC cell lines (**Figure 3**). When assessing blocking antibodies in hAT-CM, neutralization of each individual adipokine failed to suppress CXCL5 secretion. A reduction in CXCL5 secretion was only observed by combining both IL-1β and TNF neutralizing antibodies, which provided a significant suppressive effect against the pooled hAT-CM (Figure 3).

**Figure 3:**
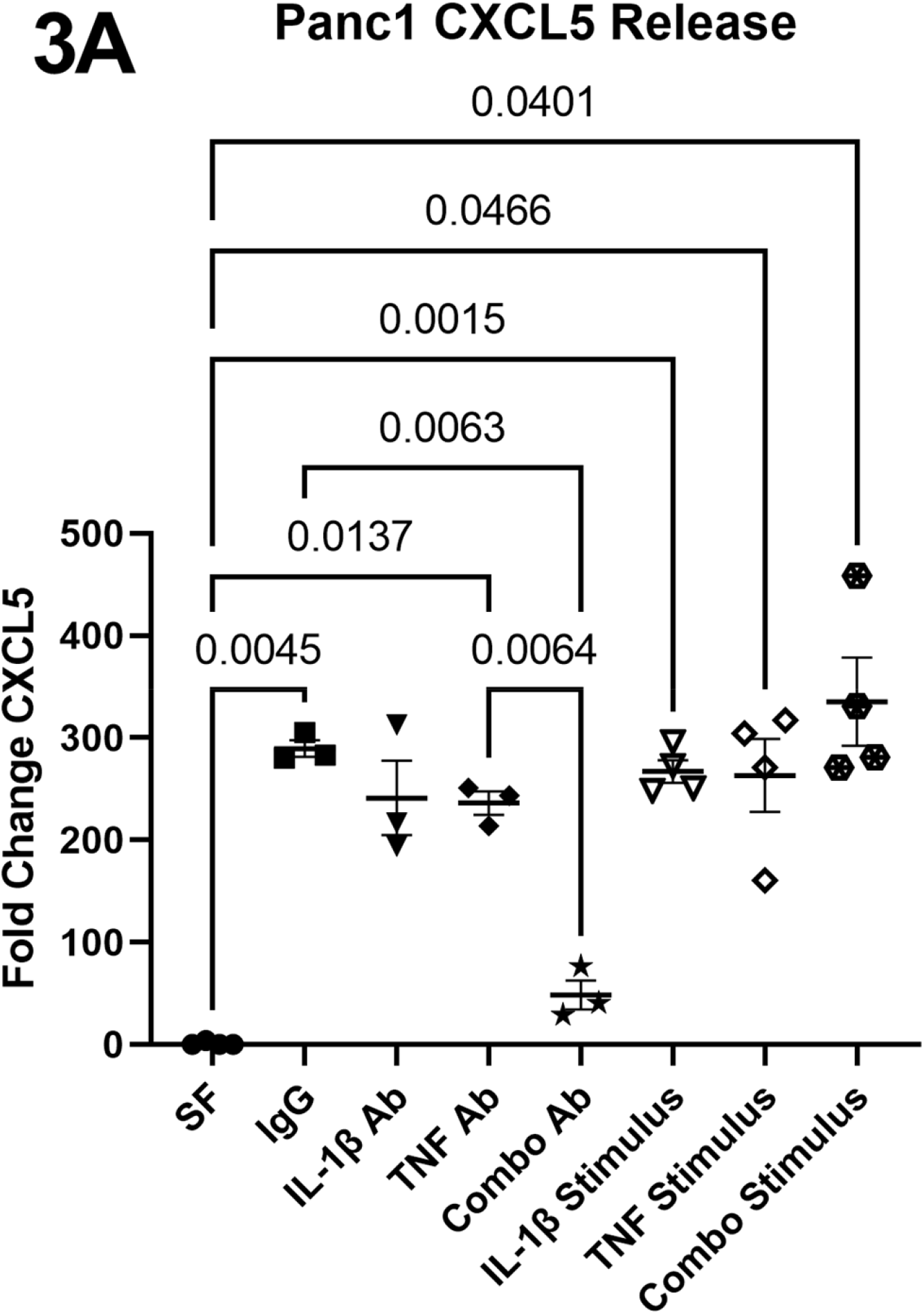
hAT-CM of pooled obese samples or recombinant IL-1b, TNF, or combo (1 ug/mL each) was used to stimulate CXCL5 release. Neutralizing antibodies against IL-1b or TNF were applied at 1 ug/mL to inhibit CXCL5 secretion in the presence of hAT-CM. Cultures were treated for 36h. Statistics were performed by Brown-Forsythe & Welch’s One Way ANOVA.

### CXCL5 deficient PDAC shows non-differential growth, but affects tumor associated myeloid profiles

In order to assess the contribution of CXCL5 on PDAC tumor progression, we sought to genetically deplete CXCL5 by CRISPR-Cas9 editing of PDAC cells. Murine K8484 PDAC cells were transfected with lentiviral particles for Cas9 expression and then subsequently with lentiviral particles for multiple CXCL5 guide RNAs (**Figure 4A**). Positive clones from each guide RNA were identified for efficient knockout by qPCR analysis (**Figure 4A**). The controls and knockouts were tested for changes in proliferation by EdU incorporation *in vitro*. We found no deleterious effect of the knockout toward intrinsic cancer cell proliferation when pooling clones of the same guide-RNA (**Figure 5B**), however, there was a significant reduction in EdU incorporation of clone 3-17, respectively, when separately compared to the control line (Supplemental **Figure 4**). To overcome potential differences between clones, two pairs of knockout clones from each guide RNA were mixed and orthotopically injected into obese mouse pancreas (**Figure 4C**). Tumor volume at endpoint failed to reveal an effect of CXCL5 knockout on growth (**Figure 4D**). Because previous studies implicated a canonical effect of CXCL5 in promoting myeloid migration as a CXCR2 ligand (23,39), we checked for differences in myeloid immune infiltration by flow cytometry. We observed no difference in gross myeloid (CD11b+) immune infiltration (**Figure 4E)**, but we did identify an insignificant trend toward increased number of antigen presenting cells (MHCII+,CD206-/CD11b+) (**Figure 4F)**. Contrary to the anticipated results, we observed a drastic increase in the presumed monocytic MDSC (Ly6G-,Ly6Chi/CD11b+) population in our CXCL5 deficient tumors (**Figure 4G)**. Additionally, using CXCR4 as a marker for pro-tumor neutrophils (40), we observed an enhancement of the polarization of the neutrophils to a pro-tumorigenic phenotype (**Figure 4H-I)**.

**Figure 4:**
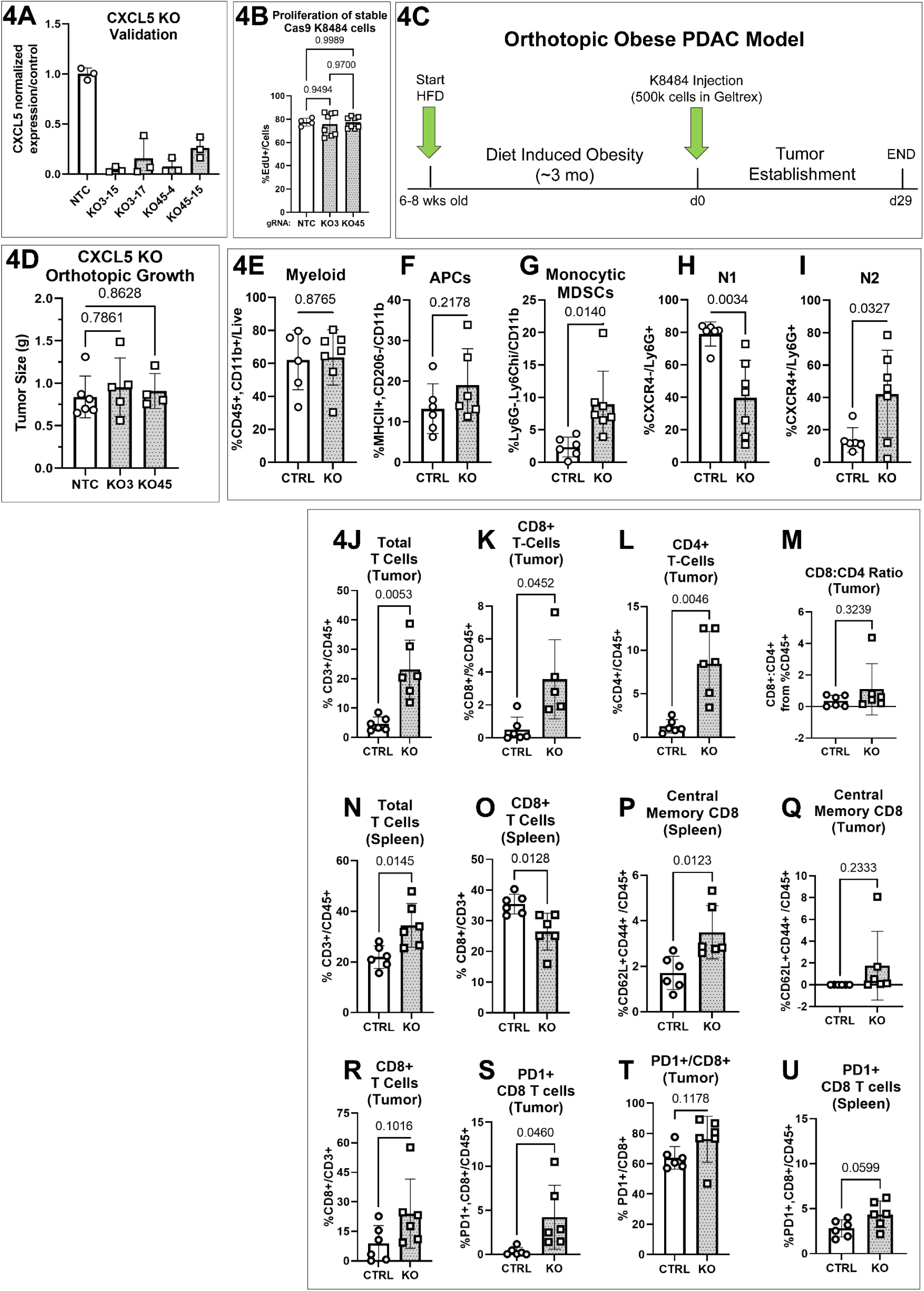
(A) CXCL5 was validated for knockout in murine PDAC cell lines (KPC) by qPCR. Proliferation of the cells *in vitro* was measured by EdU incorporation over a 6h time period in 5% DMEM. (C-U) The knockout cells were pooled (3-15 + 3-17, or 45-4+45-15) and orthotopically injected into the pancreas of obese mice. Tumors were allowed to grow for 29 days prior to takedown when they were assessed for tumor weight and immune infiltration by flow cytometry (see Supplement 4 for gating strategy and antibodies). Statistics were performed by Brown-Forsythe & Welch’s One Way ANOVA (4B, 4D), or Welch’s T-test (4E-U).

**Figure 5:**
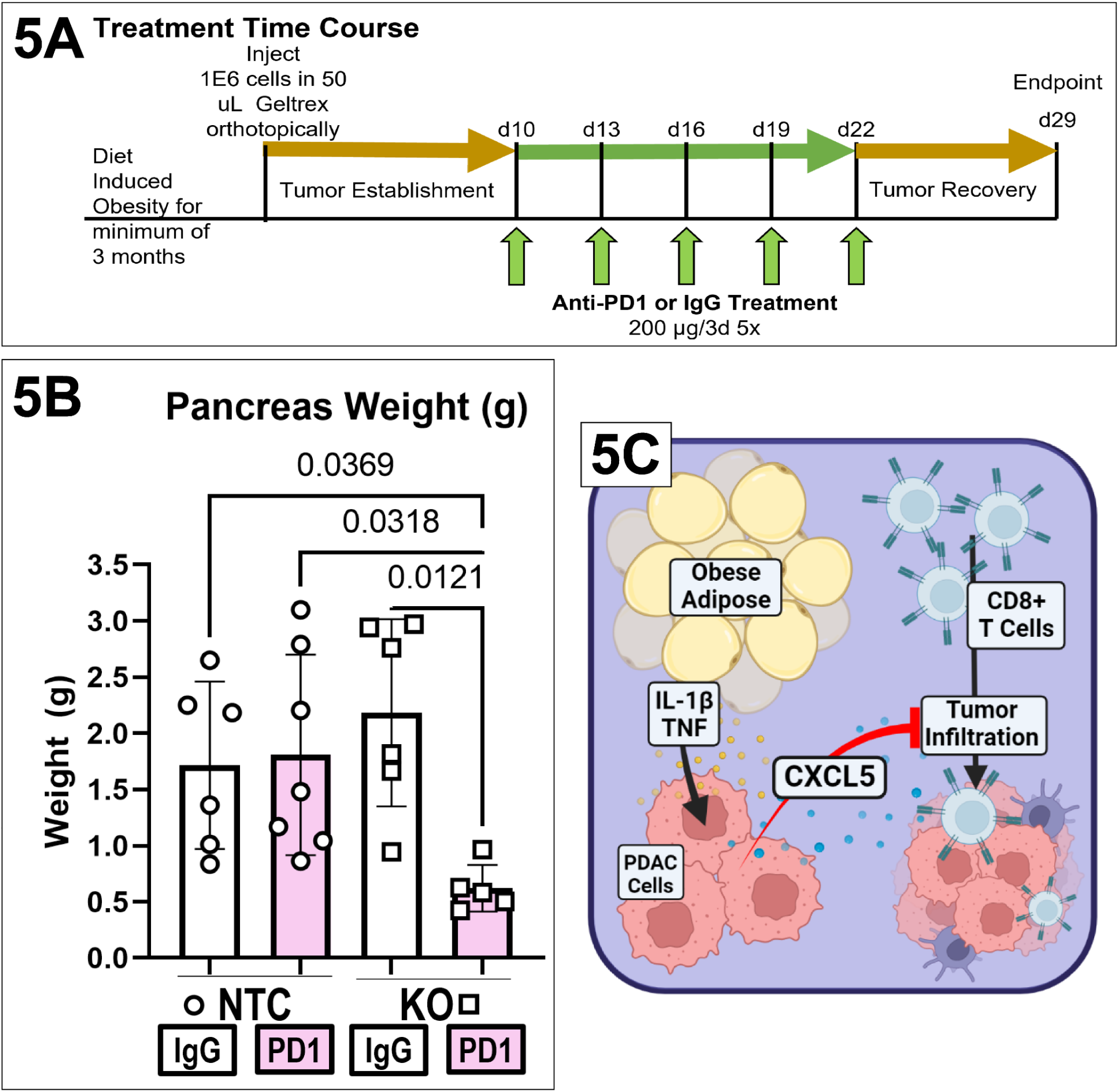
Mice underwent diet-induced obesity and tumor implantation as in Figure 4B. At day 10 post tumor implantation, mice were treated with 200 ug of anti-PD-1 antibody, or IgG and continued every 3^rd^ day until d22. Pancreas weight was assessed at endpoint (B). Statistics were performed by Brown-Forsythe & Welch’s One Way ANOVA. (C) Adipose tissue contains cytokines that can stimulate CXCL5 release from PDAC cells, which acts as a critical deterrent to tumor T cell infiltration (figure created in Biorender).

### Loss of CXCL5 enhanced CD8 T cell tumor infiltration and systemic CD8 profile

While the neutrophil and MDSC profiles of the CXCL5-knockout tumors trended toward a less favorable tumor phenotype for treatment, we noted that the T cell profile of the knockout tumors presented much more encouraging results. The CXCL5-knockout tumors revealed a drastic increase in CD3 T cell infiltration **(Figure 4J)** with increases in both CD4 and CD8 T cells (**Figure 4K-L**), yet this did not result in a significant increase in the ratio of CD8:CD4 infiltration (**Figure 4M**). Looking at the circulating T cells from the spleens of the tumor bearing mice, we found that the total T cells (CD3+/CD45+) were increased systemically in the CXCL5-KO tumor bearing mice (**Figure 4N**). Notably, central memory CD8 T cells (CD62L+,CD44+/CD8) were significantly increased in the spleens of the CXCL5-KO bearing mice, yet there was an insignificant trend in the increase in the tumor infiltrate (**Figure 4P-Q**). While the proportion of CD8:CD45 cells was increased in the CXCL5 knockout tumors, the CD8 cells as a proportion of CD3 cells did not display a significant increase (**Figure 4R**). Importantly, the CXCL5-KO tumor bearing mice CD8 immune population displayed high levels of PD-1 expression in the tumor, even though they did not have a significantly increased proportion of PD-1+/CD8+ (**Figure 4S-U**).

### CXCL5 deficient PDAC is susceptible to anti-PD1 immune checkpoint blockade therapy

The CD8+ infiltrating cells into the CXCL5-knockout tumors displayed substantial PD-1 positivity (**Figure 4S**), suggesting that combination with checkpoint inhibitor blockade therapy could provide a therapeutic benefit in a CXCL5 deficient tumor setting. NTC and CXCL5 deficient tumors were treated with an anti-PD-1 antibody therapy regimen outlined in Figure 5A. Administration of the anti-PD-1 therapy led to a significant reduction of tumor volume in mice bearing CXCL5-knockout tumors, while the same therapy provided minimal effect in our obese NTC tumors (**Figure 5B**).

## DISCUSSION

In this study, genetic ablation of tumor derived CXCL5 was accompanied with an enhanced infiltration of endogenous CD4 and CD8 T cells into the tumor microenvironment, yet the tumors failed to display evidence of benefit due to high exhaustion and suppressive immune phenotypes in the absence of tumor derived CXCL5.

IL-1β and TNF are cytokines shown to be released by adipose depots, with increased release from adipose from patients with higher BMI/obesity status reported in the literature (38), we failed to see a significant trend when comparing overweight/obese PDAC patients to leaner patients. However, the power of our study was limited by the number of patient adipose samples available, and the quantity of adipose obtained from these patients. The implications of adipokine-tumor crosstalk have been investigated intrinsically through measuring tumor proliferation, metabolism, and migration, but we observed that these adipokines can further induce the secretion of the immune modulating chemokine CXCL5 from PDAC cells.

In these studies, we observed that TNF or IL-1β are sufficient to drive CXCL5 secretion - where TNF is likely promoting CXCL5 expression via CREB, which has been reported to be sourced from tumor associated neutrophils (18). Additionally, IL-1β and TNF both contributed to CXCL5 secretion via the NF-kB pathway in inflammatory contexts (39,41). Our results show that presence of either TNF or IL-1β appears to be sufficient to promote CXCL5 release from the tumor cells, but only blockade of both limited secretion of CXCL5 in our *in vitro* approach (**Figure 3**). Notably, fatty acids from adipose, such as palmitate that was shown to be sufficient to induce CXCR1/2 ligand secretion from hepatocellular carcinoma cells (42), could be playing a role in the residual secretion after combined blockage of IL-1β and TNF in our assay.

While our analysis of TCGA data shows that CXCL5 is the only CXCR1/2 ligand that is significantly associated with survival, Steele et al. showed in analyses of biopsies from resected tumors that CXCL2 is associated with worsened patient outcome(23). CXCR2 ligands, including CXCL5, have been previously implicated in enhancing pro-tumor myeloid immune infiltration into the PDAC TME (18,23,26,39,43). Despite this canonical understanding of CXCR1/2 ligands in myeloid trafficking, alternatively, our flow cytometric analysis of CXCL5-KO tumors displayed an unexpected immune profile with an enrichment of (Ly6Chi,Ly6G-/CD11b+) cells which resemble monocytic MDSCs (**Figure 4G)**. This suggests to us that there may be differential effects of related CXCL ligands, with specific subtleties in their targets and effects:

While CXCL1 has been implicated as a critical regulator of T cell infiltration through TNF signaling (18), our data suggests that CXCL5 loss specifically leads to a tumor suppressive myeloid profile in contrast to CXCL1 loss, with the caveat that the obese TME may also be a contributing factor to the discrepancy. To elicit an anti-tumor effect, CXCL5 depletion requires checkpoint inhibition for an anti-tumor response in a manner similar to IL-8 (18,27). In contrast, CXCR2 ligand signaling was shown to affect proliferation both negatively and positively in the following studies: Application of recombinant CXCL1 *in vitro* to immortalized ductal cells induced senescence (44), yet, CXCR2 knockdown was shown to inhibit PDAC cell growth (45), and CXCL1 autocrine signaling was shown to promote proliferation of esophageal cancer cells (46). Our analysis of CXCL5 knockout cells displayed no observed effect of the loss of endogenous CXCL5 on tumor cell proliferation *in vitro* amongst three of the four clonal populations (**Supplemental Figure 4A)**, nor did we see differential tumor growth *in vivo* **(Figure 4D)**.

Our studies have uncovered potential areas for further consideration. Interestingly, CXCL1 and CXCL5 have been shown to interact by immunoligand blotting (47), and are predicted to interact *in silico* (48) **(Supplemental Figure 5)** in a heterodimer that appears similar to the homodimer state of CXCL5 in solution (49): perhaps the loss of CXCL5 in our studies is leading to a reduction in CXCL1/5 interaction that potentiates CXCL1 mediated chemotaxis of the monocytic MDSCs into the tumor, yet the function of the dimerizations need to be investigated further. Alternatively, a potential contribution of the adipokines to promote MAPK activation in an obese setting was noted after assessment of our RNAseq analysis, which could also contribute to the PD-1/PD-L1 axis, supplemental to dependence on the myeloid population as previously described (14).

In conclusion, our studies highlight the role for tumor derived CXCL5 in regulating an anti-tumor immune response. While the exact mechanism is still unclear, our results suggest that depletion of CXCL5 in the setting of obesity is sufficient to induce CD8 T cell infiltration into pancreatic tumors. Loss of tumor derived CXCL5 also increases immune suppression via increased myeloid cell recruitment and polarization. Therefore, depletion of CXCL5 requires anti-PD-1 therapy to achieve suppression of pancreatic cancer growth. While CXCR1/2 inhibitors are under clinical investigation, it has been reported that CXCR2 knockout mice are more susceptible to lung infections (50). Our current studies provide a potential alternative treatment strategy that may subvert side effects of broad spectrum CXCR1/2 inhibitors by targeting individual CXC ligands.

## Acknowledgements

This work was supported by grant R01 CA231052 from the NCI to Dr. VanSaun.

Research reported in this publication was supported by the National Cancer Institute Cancer Center Support Grant P30 CA168524 (PI: Jensen) CCSG pilot funding for Dr. VanSaun.

We thank the University of Kansas Medical Center – Genomics Core for generating the sequence data sets. The Genomics Core is supported by the Kansas Intellectual and Developmental Disabilities Research Center (NIH U54 HD 090216), the Molecular Regulation of Cell Development and Differentiation – COBRE (P30 GM122731), The NIH S10 High End Instrumentation Grant (NIH S10OD021743) and the Frontiers CTSA Grant (UL1TR002366).

The HPC used for RNAseq data analysis was supported by the National Cancer Institute (NCI) Cancer Center Support Grant P30 CA168524; the Kansas IDeA Network of Biomedical Research Excellence Bioinformatics Core, supported by the National Institute of General Medical Science award P20 GM103418; the Kansas Institute for Precision Medicine COBRE, supported by the National Institute of General Medical Science award P20 GM130423.

We acknowledge support from the University of Kansas (KU) Cancer Center’s Biospecimen Repository Core Facility (BRCF) staff for helping obtain human specimens. The authors also acknowledge support from the KU Cancer Center’s Cancer Center Support Grant (P30 CA168524)

We acknowledge the Flow Cytometry Core Laboratory, which is sponsored, in part, by the NIH/NIGMS COBRE grant P30 GM103326 and the NIH/NCI Cancer Center grant P30 CA168524.

We’d like to specifically thank key personnel Rich Hastings and Mary Markiewicz from the KUMC Flow Cytometry Core, and Alex Webster from the BRCF.

The K8484 cell line was a gift from the Dave Tuveson lab characterized in Olive et al. 2009 (28).

This work was presented at the Midwest TME Meeting 2022 and AACR Special Conference: Pancreatic Cancer 2022 (C060 Adipose-tumor crosstalk alters tumor immune profile by promoting PDAC CXCL5 secretion).

## Notes

### Competing Interest Statement

The authors have declared no competing interest.

