## Supplemental files for "Adipose-Tumor Crosstalk contributes to CXCL5 Mediated Immune Evasion in PDAC"

### Supplemental Figure 1

First plot represents the total mRNA count per sample (library size). The red horizontal line indicates the mean line.

The second plot represents the boxplot of the gene expression counts in log scale.

Barplot indicates slight differences in batches.

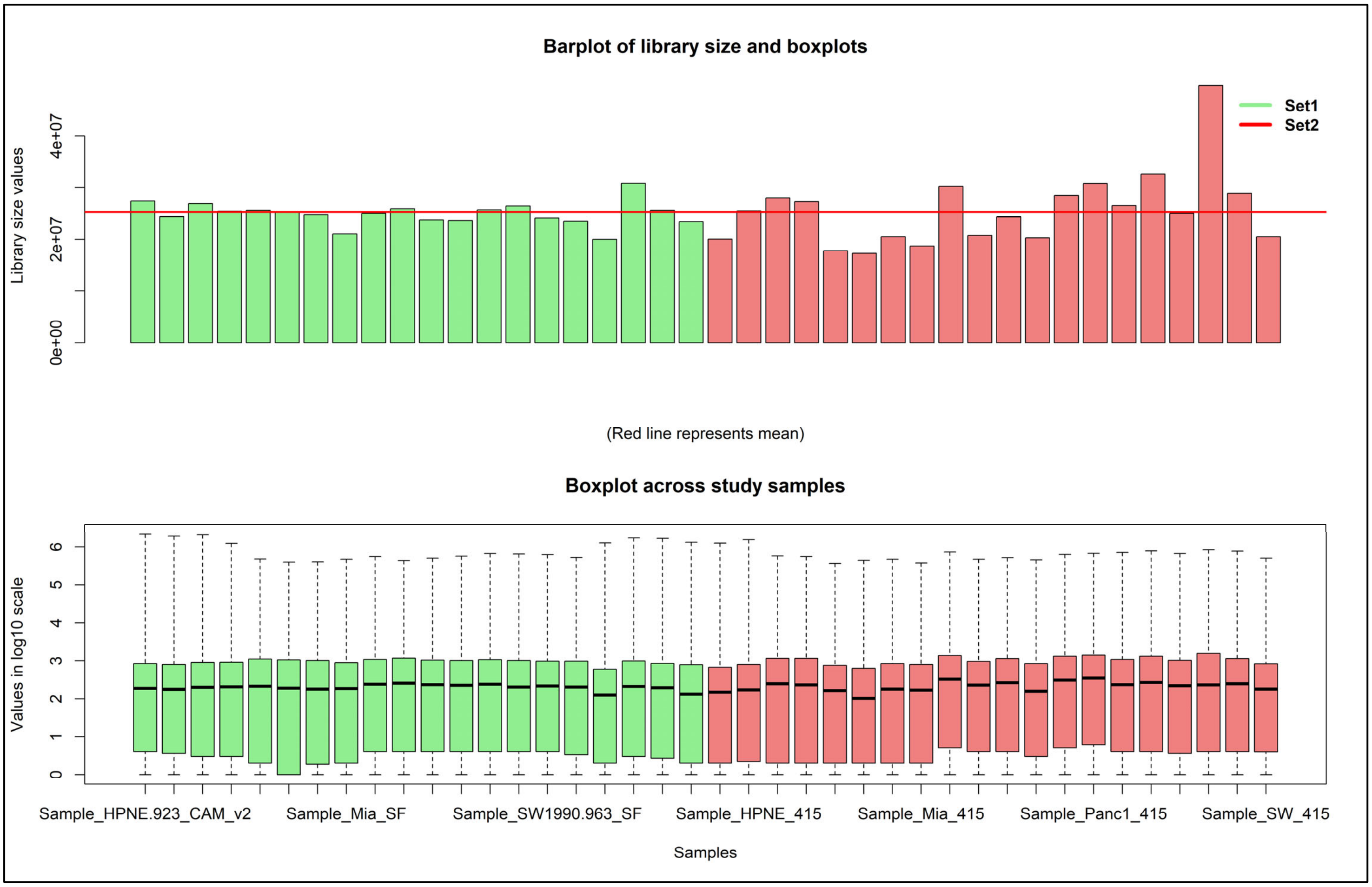

### Supplemental Figure 2

(A) Differential overall survival based on expression of CXCR1/2 ligands from PAAD data and associated p-values (A) and Kaplan-Meier Plots (B). (C) MiaPaCa2 cells stimulated with hAT-CM release CXCL5 on cytokine array. (D) IL-1b and TNF measured by ELISA from hAT-CM of lean or overweight/obese patients with PDAC. Statistics were performed by Welch's T-Test, p<.05 = *

## S2A

| Gene | CXCL1 | CXCL2 | CXCL3 | CXCL5 | CXCL7 | CXCL8 |
|---|---|---|---|---|---|---|
| Logrank p= | 0.67 | 0.13 | 0.78 | 0.024 | 0.83 | 0.59 |

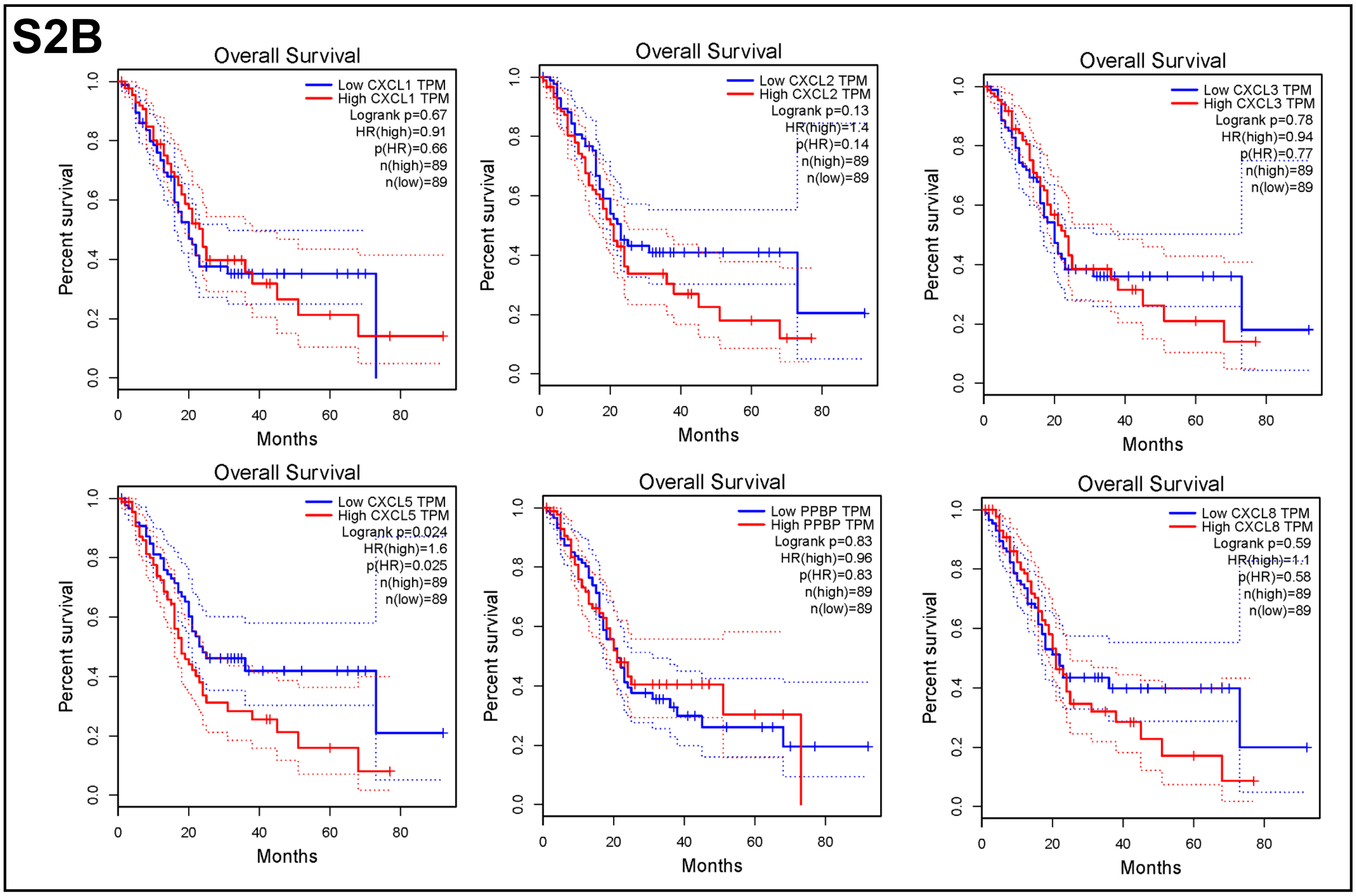

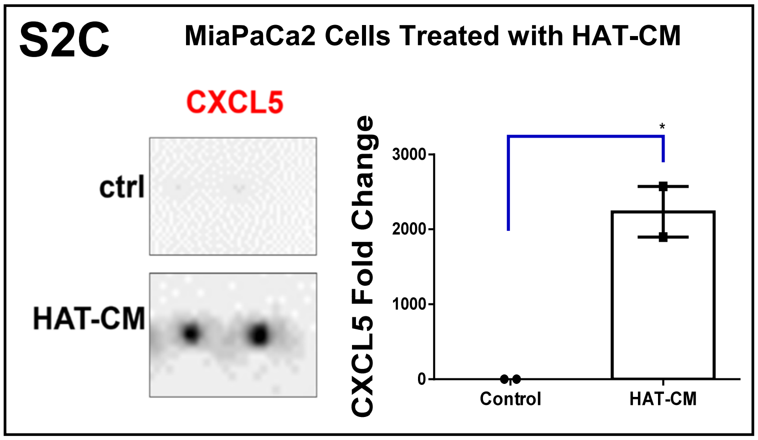

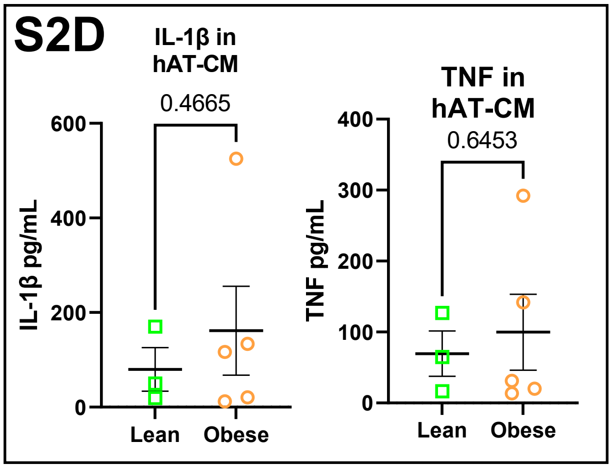

### Supplemental Figure 3

Activation of IFN pathways in IPA analysis in HPNE (A), MiaPaCa2 (B), Panc1 (C), and SW1990 (D) cells in response to Obese vs Lean hAT-CM

## S3A

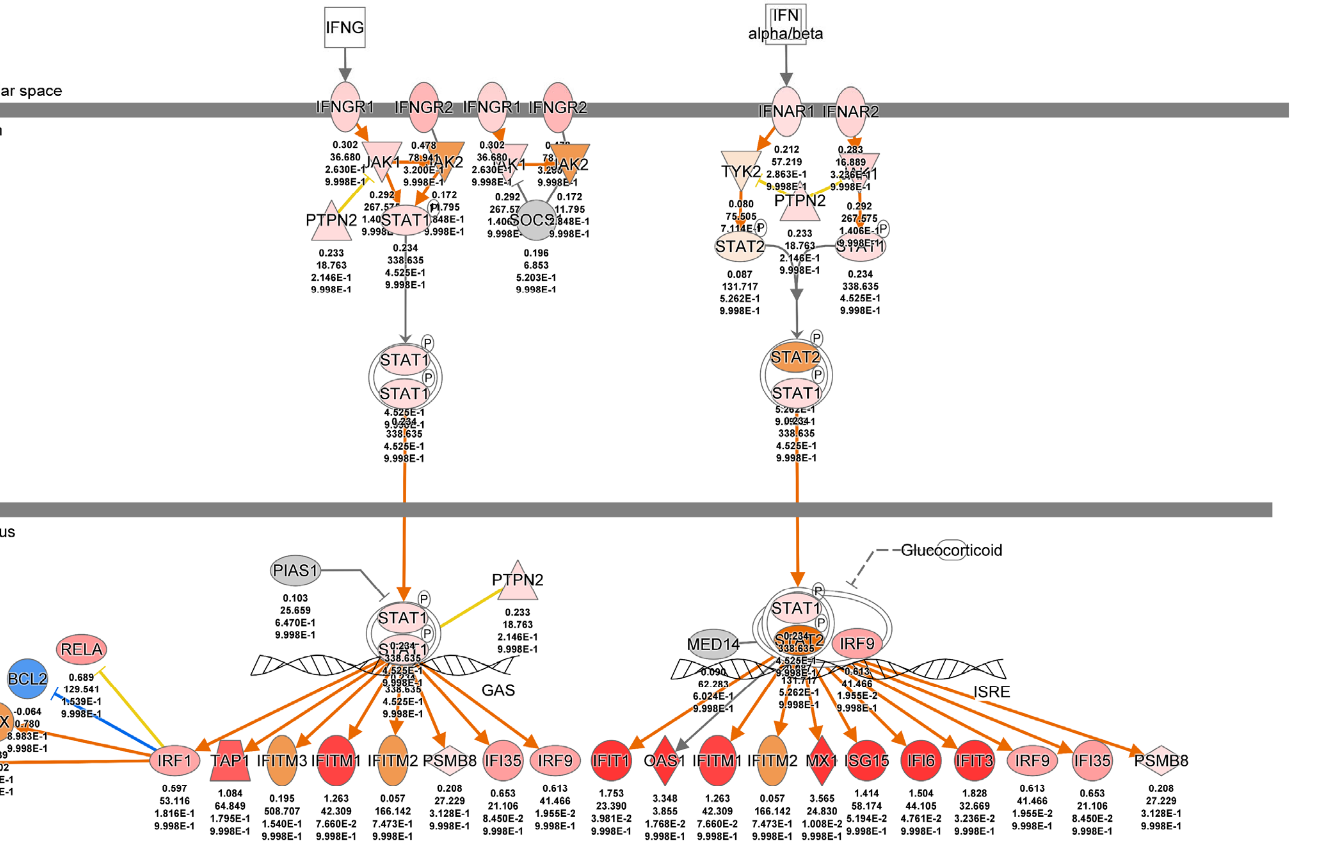

© 2000-2023 QIAGEN. All rights reserved.

## S3B

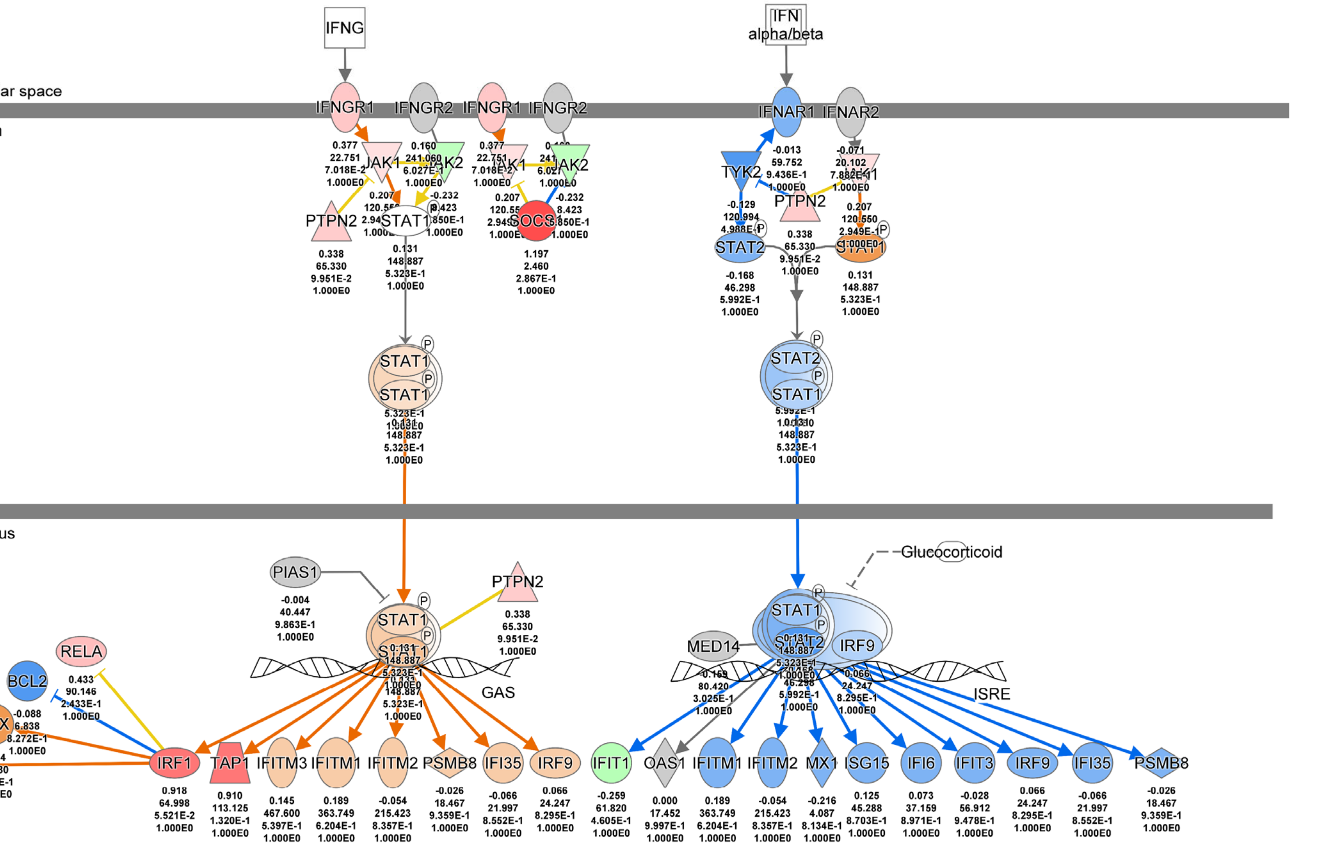

© 2000-2023 QIAGEN. All rights reserved.

### Supplemental Figure 3 (cont.)

S3C

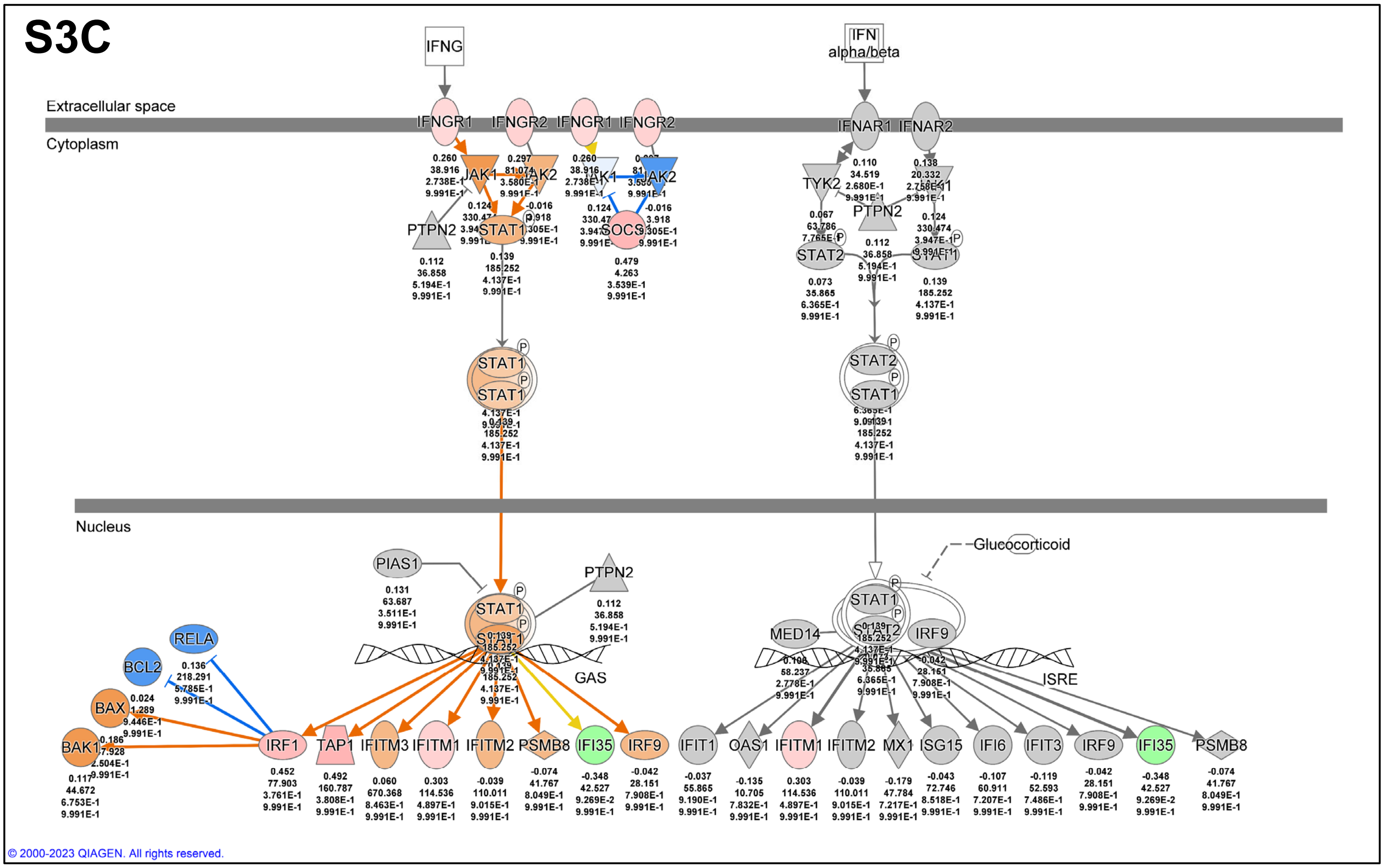

S3D

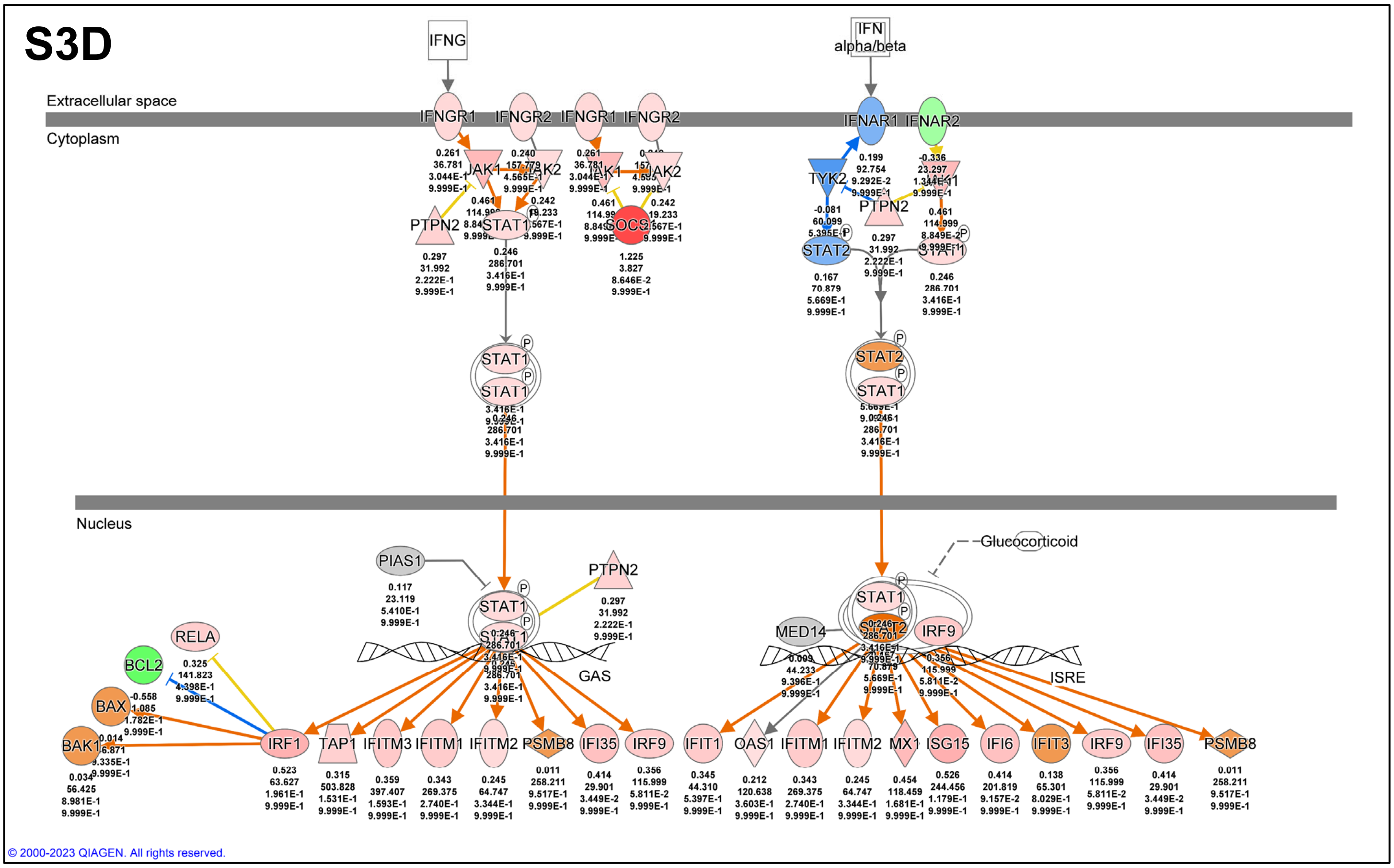

### Supplemental Figure 4:

(A) One of four clones displays reduced proliferation by measure of EdU compared to NTC (Brown-Forsythe and Welch corrected One-Way Anova). (B) Flow cytometry antibody panels. (C) Gating strategy for T cells. (D) Gating strategy for APCs. (E) Gating strategy for MDSCs and Neutrophils

## S4A

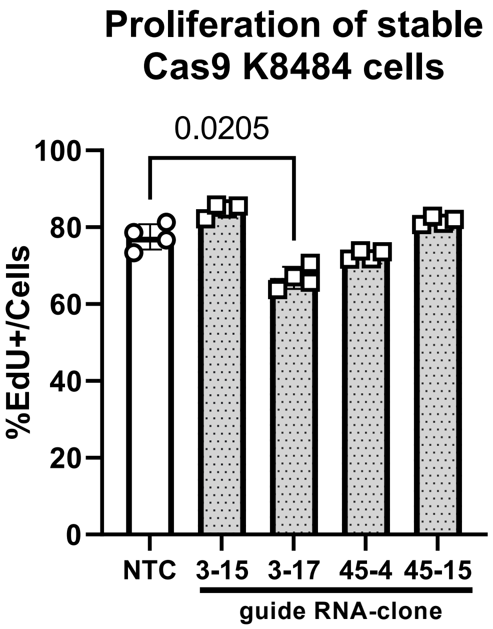

## S4B

**T Panel**

| | | | | |
|---|---|---|---|---|
| Live Dead | Aqua | ThermoFisher | N/A | L34966 |
| CD44 | BV650 | Biolegend | IM7 | 103049 |
| PD1 | BV785 | Biolegend | 29F.1A12 | 135225 |
| CD62L | FITC | Biolegend | MEL-14 | 104406 |
| CD8a (53-6.7) | PerCP Cy5.5 | Biolegend | 53-6.7 | 100734 |
| CD3e | APC | Biolegend | 145-2C11 | 100312 |
| CD4 | AF700 | Biolegend | GK1.5 | 100430 |
| CD45 | PE-Cy7 | Biolegend | 30-F11 | 103114 |
| **M1vM2 Panel** | | | | |
| CD86 | BV421 | Biolegend | | |
| CD80 | BV605 | Biolegend | 16-10A1 | 104729 |
| MHCII | BV785 | Biolegend | M5/114.15.2 | 107645 |
| CD204 | PE | invitrogen | M204PA | 12-2046-82 |
| CD206 | PE-Cy7 | Biolegend | C068C2 | 141720 |
| F480 | APC | Biolegend | BM8 | 123116 |
| CD11b | FITC | Biolegend | M1/70 | 101206 |
| **N1vN2 Panel** | | | | |
| CD25 | BV421 | Biolegend | PC61 | 102035 |
| Ly6G | BV785 | Biolegend | 1A8 | 127645 |
| ICAM-1 | FITC | Biolegend | YN1/1.7.4 | 116105 |
| Ly6C | PerCP-Cy5.5 | Biolegend | HK1.4 | 128012 |
| cxcr4 | PE | Biolegend | L276F12 | 146505 |
| CD45 | PE-Cy7 | Biolegend | 30-F11 | 103114 |
| cd135 | APC | Biolegend | A2F10 | 135309 |
| CD11b | 605nc | eBioscience | M1/70 | IH93-0112 |

### Supplemental Figure 4 (cont.)

**S4C**

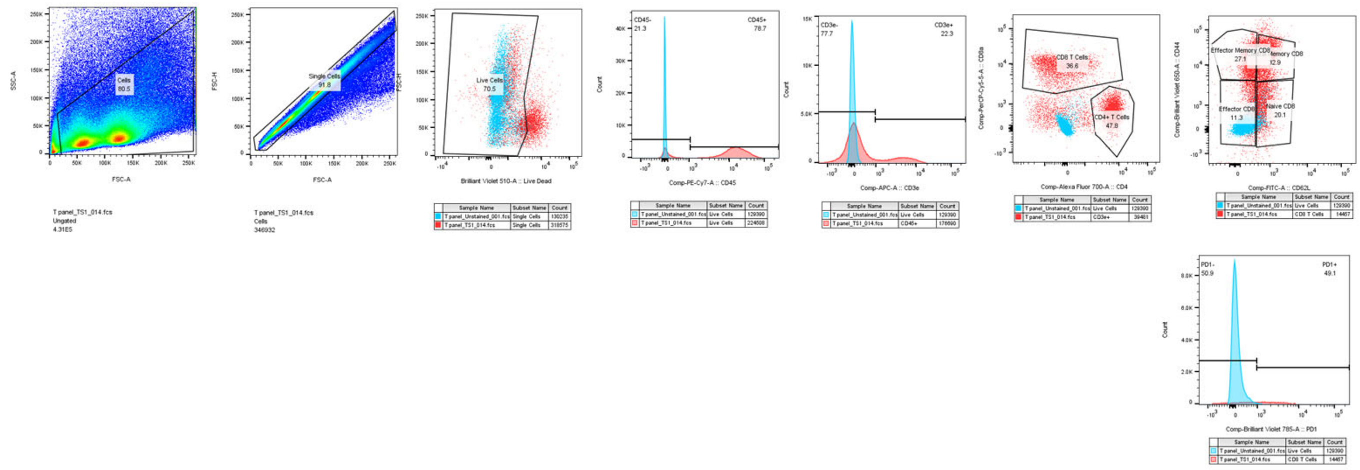

**S4D**

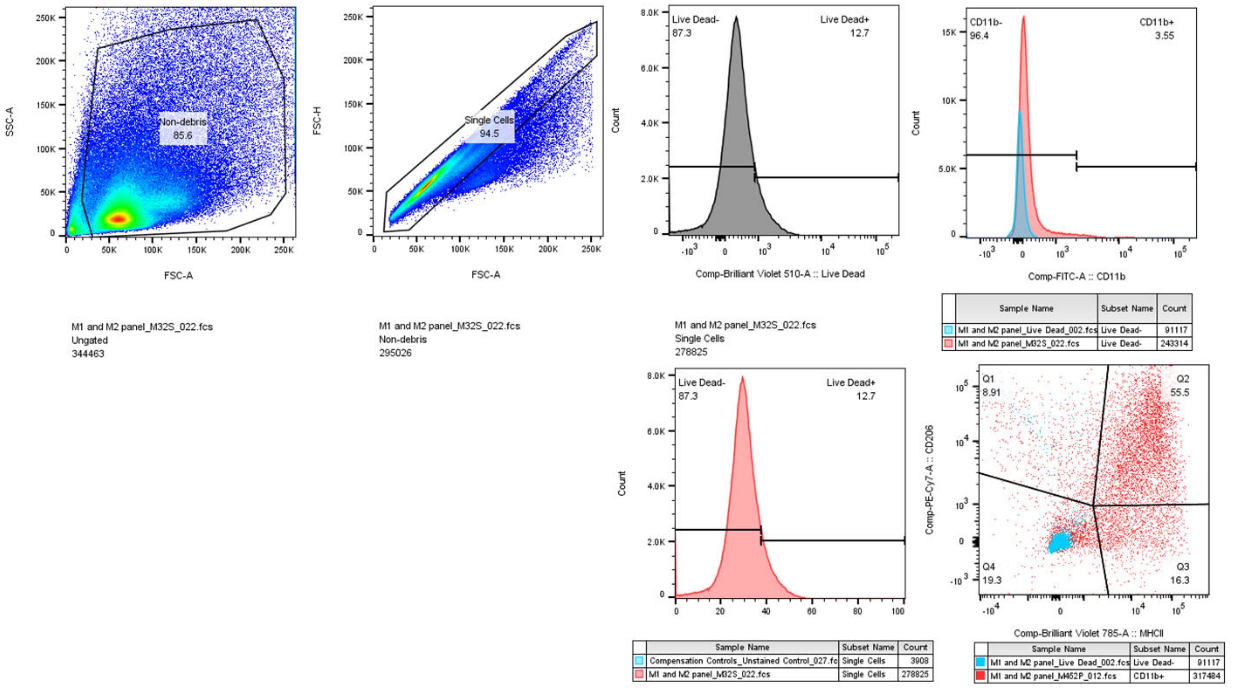

**S4E**

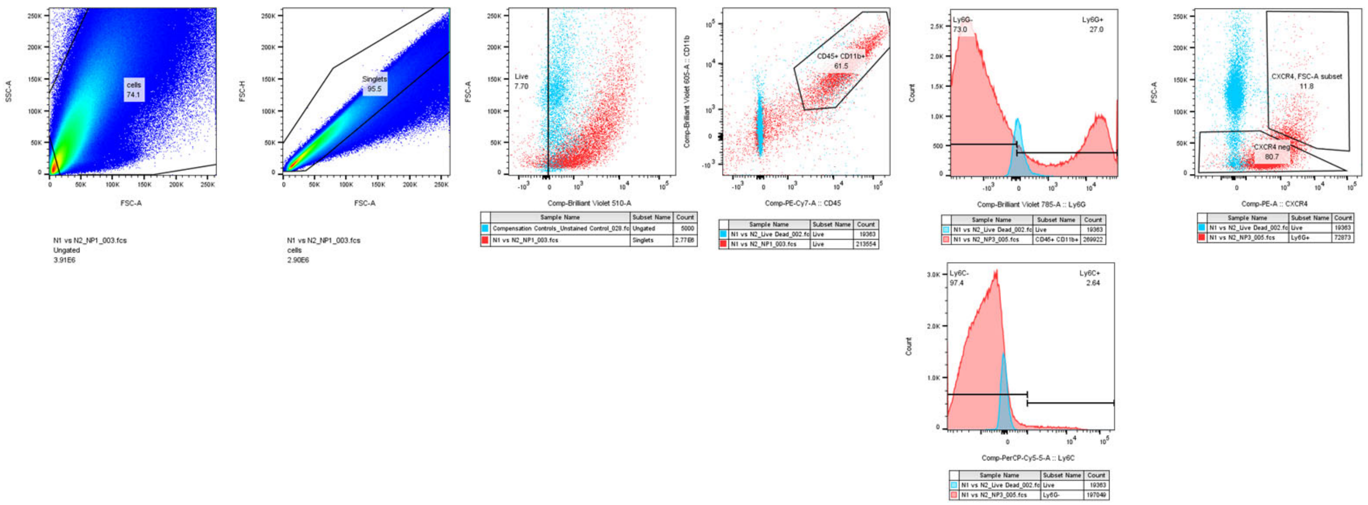

#### Supplemental Figure 5:

*in silico* analysis of CXCL5/1 binding as heterodimers using PEPPI (48) in human (left) and mouse (right) secreted versions of the ligands.

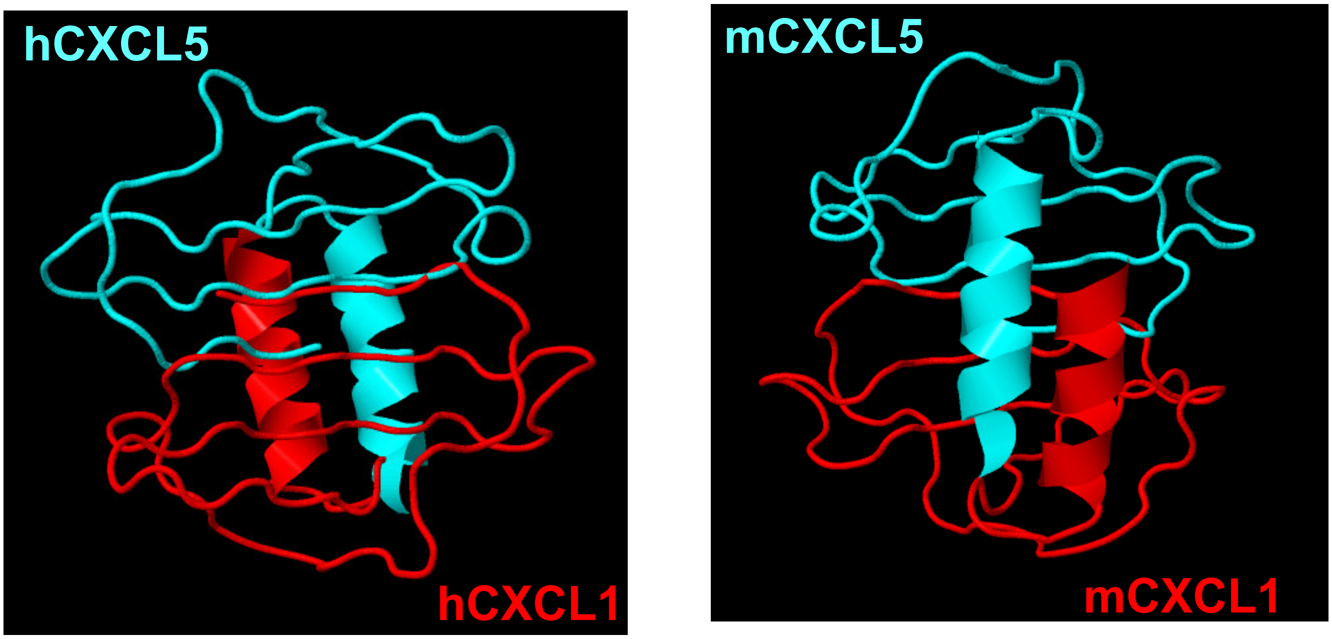

SPRING score: 17.975

SPRING score: 17.468
